# Autometa 2: A versatile tool for recovering genomes from highly-complex metagenomic communities

**DOI:** 10.1101/2023.09.01.555939

**Authors:** Evan R. Rees, Siddharth Uppal, Chase M. Clark, Andrew J. Lail, Samantha C. Waterworth, Shane D. Roesemann, Kyle A. Wolf, Jason C. Kwan

## Abstract

In 2019, we developed Autometa, an automated binning pipeline that is able to effectively recover metagenome-assembled genomes from complex environmental and non-model host-associated microbial communities. Autometa has gained widespread use in a variety of environments and has been applied in multiple research projects. However, the genome-binning workflow was at times overly complex and computationally demanding. As a consequence of Autometa’s diverse application, non-technical and technical researchers alike have noted its burdensome installation and inefficient as well as error-prone processes. Moreover its taxon-binning and genome-binning behaviors have remained obscure. For these reasons we set out to improve its accessibility, efficiency and efficacy to further enable the research community during their exploration of Earth’s environments. The highly augmented Autometa 2 release, which we present here, has vastly simplified installation, a graphical user interface and a refactored workflow for transparency and reproducibility. Furthermore, we conducted a parameter sweep on standardized community datasets to show that it is possible for Autometa to achieve better performance than any other binning pipeline, as judged by Adjusted Rand Index. Improvements in Autometa 2 enhance its accessibility for non-bioinformatic oriented researchers, scalability for large-scale and highly-complex samples and interpretation of recovered microbial communities.

**Graphical abstract:** Autometa: An automated taxon binning and genome binning workflow for single sample resolution of metagenomic communities.

## INTRODUCTION

Metagenomics enables the study of organisms that have thus far eluded cultivation, by negating the need for the isolation of pure strains or indeed any laboratory culture prior to sequencing (1). Such direct environmental sequencing and subsequent assembly generally yields contigs from a complex mixture of species, and the *de novo* separation of contigs into individual genomes (“binning”) remains a computational challenge (2). We previously developed Autometa, an automated binning pipeline that is able to effectively recover genomes from highly convoluted environmental and non-model host-associated microbial communities (3). This tool has seen widespread use in environments ranging from marine, freshwater and terrestrial samples, including corals (4), red algae (5), kinetoplastids (6), deep sea geothermal vents (7–9), sponges (10–12), coastal sediments (13), stromatolites (14), seaweeds (15), shipworms (16), plateau lakes (17, 18), hot springs (19, 20), contaminated rivers (21), beetles (22–25), Kickxellomycotina fungi (26), Ensifera insects (27), fermented agave (28, 29), a marsh orchid rhizobiome (30), domesticated cattle (31, 32), mice (33) and human gut (34), periodontal (35) as well as urinary tract (36) microbiomes. As a consequence of Autometa’s widespread use, both non-technical and technical researchers alike have communicated their frustrations regarding the ease of installation as well as the efficiency and robustness throughout the various stages of the Autometa workflow. Moreover, the behaviors of Autometa’s taxon-binning and genome-binning processes have remained obscure. We originally envisioned Autometa’s taxon classification process simply as an aid to genome binning in complex samples, and therefore the performance of taxon classification has not been rigorously benchmarked (3). Likewise, while a number of parameters are user-configurable in Autometa, we have previously not systematically explored their effects on binning performance. On account of Autometa’s diverse applications and increasing user base, we set out to address and improve upon these issues. Here we present the highly augmented Autometa 2 release. Autometa 2 comes with many enhancements in performance, maintainability and accessibility. This includes new features such as additional parameters regarding pre-processing, taxon binning and genome binning, version-controlled documentation, tooling for continuous integration, testing, benchmarking and deployment, and finally modularization which provides all of these metagenomics processing features through a python API.

Additionally, automated recovery of high-quality genomes from highly-complex samples or samples with high degrees of micro-diversity remains recalcitrant, largely because of the required time and compute requirements. Due to the size of these ever-increasing metagenomic datasets, we created a “large data mode” and benchmarked the computational requirements and performance metrics for both Autometa versions 1 and 2, as well as “large data mode”, alongside other common metagenomics software. We characterized Autometa’s genome binning performance and examined binning behavioral changes based on 33, 158 parameter configurations using the CAMI 2 datasets (2), and provide additional descriptions of its taxon binning approaches. With these binning insights we outline next steps in developing methods to accommodate datasets deemed intractable due to complexity and scale. Non-model host-associated samples often fall under this category and are of particular interest to the Kwan Group (10, 14, 22, 37). We also highlight the inherent difficulties with recovery of highly-resolved genomes from highly-complex and host-associated metagenomes.

## MATERIAL AND METHODS

### Workflow overview

The workflow for Autometa 2 is largely unchanged from Autometa 1, with the exception that there are now more parameters that users can control. Briefly, Autometa first performs pre-processing tasks where assembled contiguous sequences (contigs) are filtered by length and taxon. The latter process assigns contigs to kingdom-level taxonomies, effectively separating eukaryotic host-associated genomes from prokaryotic symbionts. Contigs are recursively binned using nucleotide composition and read coverage, with successive rounds first splitting the remaining contigs into groups from less to more specific canonical ranks (i.e. kingdom, phylum, class, order, family, genus, species). Finally, Autometa attempts to recruit any remaining unclustered sequences into one of the recovered putative metagenome-assembled genomes (MAGs) through classification by a decision tree classifier (or optionally, a random forest classifier). The resulting MAGs may then be subjected to manual inspection prior to downstream comparative genomic analyses (a companion graphical interface, Automappa (38), was developed specifically for this and may be found at: https://github.com/WiscEvan/Automappa).

The general Autometa workflow (Figure 1) consists of eight stages:

1. Length-filtering: Discard short sequences that may resemble assembly artifacts
2. Coverage analysis: Calculate contig coverage (a proxy for abundance)
3. *K*-mer analysis: Determine sequence similarity based on nucleotide composition
4. ORF-calling: Identify contigs’ open-reading frames (ORFs)
5. Marker-annotation: Identify and annotate single-copy marker genes
6. Taxon-binning: Assign and group contigs by predicted taxonomic rank
7. Genome-binning: Single-copy marker gene guided recursive clustering of contigs into MAGs
8. Unclustered recruitment: Using aggregate features of the recovered MAGs from genome-binning, attempt to recruit unclustered contigs into their corresponding MAG

**Figure 1.**
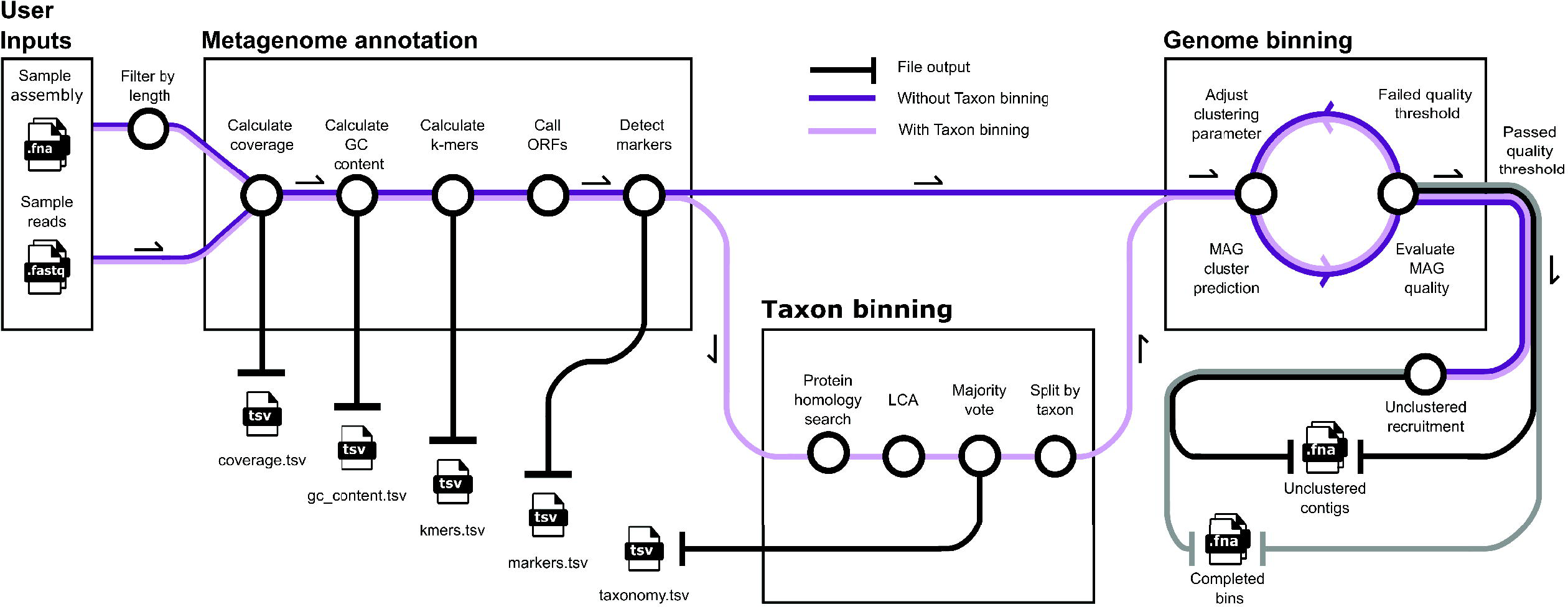
An overview of the Autometa workflow, depicting two possible routes. The first route performs genome binning without further metagenome processing. The second route incorporates Autometa’s taxon binning subworkflow as an additional metagenome processing task. Routes depicted are parallelized where possible when using Autometa’s nextflow workflow. Multiple user inputs may be provided with an input sample sheet for concurrent processing (and checkpointing) of multiple metagenomes.

### Modularized code/workflow

The Autometa library has been refactored into individual modules with submodules, using an object-oriented approach. This both lowers the barrier of entry for code contributions, and simultaneously provides multiple commands to process metagenomic data before, during and after the Autometa workflow. Due to the modular structure of the source code, community-requested features may be readily integrated into the suite of Autometa commands. As an example, we integrated the Genome Taxonomy Database (GTDB) (39) as an additional database that can be used during the taxon-binning and genome-binning stages, in place of the NCBI taxonomy, and similarly benchmark Autometa’s genome-binning performance with GTDB in use.

### New file and input options

The previous Autometa version had a number of file handling limitations during database and metagenome pre-processing. Database preparation required de-compressed inputs, greatly increasing the disk requirements as well as limiting the versatility of incorporating user-specific databases. Autometa 2 is now capable of handling gzipped inputs and outputs, vastly lowering the user’s disk requirements. Both the Autometa Python library and the new Nextflow workflow assist users in downloading and formatting all required databases. Additionally, many of the intermediate results throughout the workflow may now be written for user inspection, if desired.

### Version controlled documentation and tutorials

Autometa 2 is accompanied with major improvements in its documentation. This includes detailed instructions on how to install and configure Autometa to suit user-specific needs. Detailed explanations are outlined in the available walkthrough tutorials for options regarding which workflow to use, how to manage Autometa’s dependencies, how to select the appropriate parameters for running the workflow and how to interpret results. With a view to encouraging user-submitted code improvements and fixes, we have also released contributing guidelines, and welcome input from the community.

The documentation is open source and freely accessible at https://autometa.readthedocs.io. In addition to making the front-end of the pipeline easily accessible we have put considerable effort into documenting the code itself. Autometa 2 contains type hints and docstrings throughout to encourage reuse of existing functions and classes. Type hints allow developers to quickly discern the necessary inputs and outputs and this was done with the vision to make the codebase more readable and thus support contribution and feedback from the bioinformatics community. We have also adopted one of the most widely accepted style guides (Numpy) and have standardized the code-format using the uncompromising code-formatting tool, Black (40).

### Ease of installation

The restructured Autometa package is now amenable to common and easy installation options such as pip and conda. A docker image is also available, at https://hub.docker.com/r/jasonkwan/autometa, allowing the package to be shared and run in most compute environments. These Autometa distributions have also allowed widespread use through the Open Science Pool, making Autometa available to a wide array of distributed high-throughput computing facilities (41). Furthermore, Autometa is continuously deployed, automatically pushing each release to the bioconda channel for use with the conda package manager.

### Integration with a workflow management framework, nf-core

Autometa is complex and makes use of multiple bioinformatic databases, software packages and methods that can use significant cpu, disk and memory. To glue the different steps together into a comprehensive workflow, and to parallelize processes where appropriate, we created an optional Nextflow workflow using the nf-core framework (42, 43). This integration provides robustness, efficiency, reproducibility and scalability. Autometa’s Nextflow workflow may be configured to submit tasks to a local machine, lab servers, and a variety of cloud compute infrastructures. Nextflow also comes equipped with checkpointing whereby, if the Autometa workflow were to be interrupted, tasks may be resumed from their most recently completed process. Additionally, nf-core offers a browser-based submission interface allowing Autometa users to easily configure their metagenomic analyses. Utilizing this workflow management framework enables the submission of multiple metagenomes with one submission file using a single command, which previously would have required constant monitoring, possible re-submissions and a great deal of effort by the end-user in ensuring each metagenome’s successful processing.

### Benchmarking

A Github repository (https://github.com/KwanLab/MetaBenchmarks) has been provided containing this study’s benchmarking methods. This is an open-source repository allowing other users to regenerate metagenomics benchmarks for taxon binning and genome binning performance comparisons. Its primary purpose is to provide transparency and reproducibility when comparing metagenomics tools and has ultimately resulted in a repository where users may add their own tools for benchmarking comparison.

Altogether, fifteen tools were benchmarked against Autometa versions 1 and 2 using two different datasets. These datasets included assemblies which were taken from the second round of the Critical Assessment of Metagenome Interpretation challenge (CAMI 2) (2) and simulated communities which were initially published with the first Autometa release (3). Here, we concentrated on comparing binning performance using CAMI 2 datasets, while previously published simulated communities were used for taxon binning assessment.

For genome binning benchmarks, we utilized previously published results using the CAMI 2 datasets (2) to compare Autometa 2 to CONCOCT, MetaBAT2, MaxBin2, Vamb and Autometa version 2 as well as git commit “146383e” of Autometa version 1, which is the version used for the CAMI 2 challenge (44–47). Genome binning refiners were included in the CAMI 2 benchmarks and therefore MetaWRAP (48) and UltraBinner (https://github.com/huangpq2019/ultrabinner) were included here for comparison, however it should be noted these are not *de novo* genome binning algorithms, but perform consensus clustering via MAGs obtained by the aforementioned genome-binning tools. Taxonomy binning benchmarks used both CAMI 2 datasets as well as the simulated communities. Previously published results using the CAMI 2 datasets (2) were retrieved for Kraken 2.0.8-beta (49), Diamond 0.9.28 (50), LSHVec cami2 (51), MEGAN 6.15.2 (52), and PhyloPithiaS+ 1.4 (53). For the simulated communities dataset Autometa v1.0.3 and v2.1.0 as well as MMSeqs2 v13.45111 (54), Kraken 2.1.2 (49) and Diamond v0.9.21.122 (50) were compared. Taxon binners were selected according to their intended use on metagenome assemblies (e.g. contigs rather than reads), availability as an open-source command-line utility and synchronization with NCBI’s taxonomy databases.

A variety of clustering and classification metrics were utilized to assess binner behavior and performance, based on rationale offered by the CAMI 2 assessment (2). Briefly, following the example of the CAMI 2 challenge, we adopt the combined use of Adjusted Rand Index (ARI) and the percentage of binned base pairs in the dataset as a measure of binning performance. The ARI measures the number of true positive matchings of base pairs binned together and the number of true negatives binned apart as a proportion of the binned fraction of base pairs, with 1 being a perfect score and 0 being no better than chance (55). We additionally measured precision, recall and F1 score. Metrics were sample-weighted by both sequence count (seq) and sequence length (bp). MAG-related metrics of completeness, purity, binned percentage (sample-weighted as above) and percentage of genomes recovered (sample-weighted as above) were assessed using the previously described methods (3, 56).

To calculate these metrics for the CAMI 2 data we adapted the benchmarking tool Assessment of Metagenome BinnERs (AMBER) from the commands employed for Meyer *et al.* (2). For comparison against the simulated communities (that use different ground truths and databases), precision, recall and F1 scores were computed using the available Autometa command ‘autometa-benchmark’ (Figure S5). Future binning studies may easily extend MetaBenchmarks to perform custom parameter sweep analysis.

### New methods to expand taxon binning

Following the length-filter task, the second optional pre-processing step during the Autometa workflow is taxon binning. Autometa determines the lowest common ancestor (LCA) of ORFs based on results from a protein database similarity search. The resulting ORF LCA annotations are reduced by modified majority vote to assign taxonomic information to individual contigs (3). The rationale for applying a majority voting scheme to each contigs’ ORFs is to reduce the confounding impact of horizontal gene transfer. Assigning taxonomy to contigs allows for two features of Autometa - the separation of sequences from different kingdoms (for example, host and microbiome) and the partitioning of contigs into simpler subfractions in order to simplify clustering. The first version of Autometa only allowed domain-level taxon-filtering, whereas, this module is now capable of removing contigs corresponding to a user-provided taxon at a user-provided canonical rank. The default setting is to remove contigs outside of the kingdoms of bacteria and archaea. This has previously been shown to be a crucial pre-processing step particularly for host-associated metagenomes (3). Autometa uses Prodigal (57) for ORF prediction and Diamond (50) for accelerated protein sequence alignment against NCBI’s non-redundant (nr) protein database (50, 57). The updated version of Autometa performs Prodigal’s ORF prediction in parallel (using GNUParallel), thereby decreasing Autometa’s overall runtimes.

Autometa’s modularization allows the user to pick and choose different tools for each stage of the workflow. This includes homology search methods, LCA identification, voting schema and filtering of putative taxon-specific contaminants. One deviation from the previous Autometa workflow comes from a new taxonomy database integration, GTDB (39, 58). This integration accommodates the revised designations outlined within GDTB to account for many of the new and highly divergent microbes recovered from global metagenomic surveys. Another advantage of GTDB is its rationalization of canonical ranks in terms of sequence divergence, which should improve sequence-based taxonomic classification. However, one limitation of GTDB is that it only provides taxonomic designations for bacteria and archaea, therefore, a dual NCBI and GTDB approach must be taken if a metagenome is predicted to contain sequence from other kingdoms. This proceeds first with filtering out any contaminating kingdoms (i.e. Eukaryota, Viruses) using NCBI’s non-redundant protein database then subsequently assigning taxonomy for bacterial and archaeal fractions using the GTDB database. The GTDB taxon results are then used as the taxon annotations during the genome binning stage. This incurs additional overhead as two iterations of the protein database similarity search must be performed (i.e. once against NCBI and another against GTDB). However, since the GTDB database contains a combined size of ∼64 gigabases (Gb) (approximately 300, 000 prokaryotic genomes) compared to ∼163 Gb in nr, database searches are much faster during GTDB classification.

### New methods to expand genome binning

#### Sequence similarity analysis

Sequence similarity by nucleotide composition is performed using an alignment-free method that counts *k*-mers (sub-strings of *k* length) present in each contig. *K*-mer counts are normalized, subjected to principal component analysis (PCA), then reduced to two dimensions via a dimension reduction approach known as embedding. This subworkflow is commonly referred to as a *k*-mer frequency analysis and Autometa 2 contains multiple new parameters at each process of this stage. Autometa is packaged with a new entrypoint (’autometa-kmers’) that provides options such as *k*-mer length, normalization method, PCA dimensions, embedding method and embedding dimensions. Additionally, runtimes have been reduced by parallelization of the *k*-mer counting process. With these additional parameters, the user now has multiple perspectives with which to analyze their metagenome. For example, each embedding method performs dimension reduction in a different manner leading to different coordinates in the embedded space. These various embeddings may be visualized to better interpret the relationships between sequences and their corresponding MAGs. An example analysis on the CAMI 2 marine gold standard assembly dataset is shown in Figure S2.

#### Scaling to high-complexity metagenomes

Autometa’s original taxon-guided genome-binning approach recursively clusters contigs as it iterates through the metagenome’s taxon bins. This is performed in an ordered and sequential manner from kingdom to species where, following each iteration, unbinned contigs are passed on to the subsequent and more specific taxon subfractions. However, highly-complex metagenomes may contain many contigs in a corresponding taxon bin causing prohibitively large memory requirements and lengthy runtimes for the clustering process. Therefore, Autometa 2 is equipped with a new “large-data-mode” binning module (autometa-binning-ldm entrypoint) which was optimized to handle the most complex metagenomes that have been assembled to date.

The pseudocode depicting the “large-data-mode” algorithm may be found in Figure S3. Briefly, similar to the original taxon-guided approach, Autometa proceeds by sequentially iterating through taxon bins. At this stage, rather than the proceeding with the original method where the taxon bin is subjected to recursive clustering and MAG quality analysis, “large-data-mode” validates the taxon bin size according to whether it is within the user-designated taxon bin size range (e.g. the ‘--max-partition-size’ parameter). If within range, the taxon bin undergoes its own *k*-mer frequency embedding immediately prior to the aforementioned identical recursive clustering approach. If the taxon bin has been deemed too complex, i.e. greater than the user-designated taxon bin size, Autometa further partitions it (without clustering) in the next iteration. The rationale for skipping a partition that is too complex is that clustering will take an inordinate amount of time and resources and will likely not result in high quality bins. An edge case arises with the inclusion of taxon bin specific *k*-mer frequency analyses. The dimension reduction technique utilized during the *k*-mer frequency analysis requires taxon bin sizes in terms of number of contigs be greater than the user’s specified dimensions for both the initial PCA reduction step (if applicable) and the embedding step. This edge case can occur when a taxon has only a few representatives, reflecting a relatively simple taxon bin. Under these circumstances the *k*-mer coordinates corresponding to the simple taxon bin are retrieved from a pre-computed canonical rank *k*-mer frequency analysis. Following the determination of *k*-mer frequency analysis coordinates, recursive clustering proceeds in the same manner as mentioned above. The “large-data-mode” method also replaces the previously used default clustering algorithm, Density-based Spatial Clustering of Applications with Noise (DBSCAN) with its corresponding hierarchical approach (HDBSCAN) (59–61) due to its improved scalability and memory usage.

In addition, each iteration’s *k*-mer analysis and genome binning are checkpointed, to allow the pipeline to restart at the latest assessed taxon in the event of an interrupted Autometa run. These checkpoints and their accompanying outputs also allow for more granular inspection of Autometa’s binning decisions.

### Genome binning optimization by parameter sweep benchmarking

#### Classification performance evaluation

Autometa version 1 (cami2 branch commit-146383e) (3) was assessed during the second round of the Critical Assessment of Metagenome Interpretation (CAMI) challenge, which benchmarked a variety of recently published taxon and genome binning tools (2). The CAMI 2 competition used metagenomes constructed to represent different complexities currently challenging genome binning methods including a marine dataset consisting of highly divergent genomes (“marine”) and a dataset of many closely related strains (“strain madness”). The CAMI team published gold standard assemblies (GSA) as well as megahit assemblies (MA) for each of these sample types, totaling four metagenome assemblies available for binning (2). The published CAMI 2 benchmarks revealed that Autometa v1 performed higher than most tools in some metrics (average purity) and worse for others (average completeness, F1 score, ARI). The consistently low average completeness of MAGs recovered from the Autometa v1 results may have been due to the default Autometa v1 parameters used for the CAMI submissions: 20% minimum MAG completeness, and 95% minimum MAG purity (based on single-copy bacterial marker counts per MAG). It should be noted that Autometa’s default settings are set for low completion and high purity because our lab primarily focuses on host-associated bacteria which often have reduced genomes. Consequently, the metrics published by the CAMI challenge (2) closely reflected Autometa’s default settings (average completeness ranged from 2.5-32.2% and purity 91.7-94.3%). This suggested that increasing the completeness threshold within Autometa’s own selection criteria may increase performance across these communities.

To explore this, the raw CAMI 2 benchmarks for Autometa 1 and other binning pipeline reported in the original paper were retrieved and computed alongside Autometa’s new binning methods and sweeping a selection of run parameters now possible with Autometa 2’s user-controlled settings (Table 1). Binners were assessed using a variety of metrics including: average MAG purity, completeness, F1 score and adjusted Rand index (ARI), all used in the CAMI 2 publication (2). Metrics were calculated using AMBER v2.0.3, a tool produced for the CAMI 2 challenge (56). The Autometa parameters swept included clustering methods, as well as four cutoffs applied during the genome binning process. The cutoff parameters correspond to four MAG properties which ultimately determine whether to retain the current MAG prediction during binning: single-copy marker completeness and purity as well as coverage standard deviation and GC content standard deviation. Cutoffs for MAG completeness and purity were assessed from 10% to 100% in increments of 10%. GC content and coverage standard deviations were assessed at values of 2, 5, 10 and 15. The sweep totaled 1, 296 parameter combinations per clustering method, resulting in 2, 592 configurations per sample (10, 368 total jobs from all four CAMI 2 datasets). Large-data-mode takes additional parameters for *k*-mer methods such as normalization and embedding. Two normalization methods were selected: center log-ratio and isometric log-ratio transform. Two embedding methods were selected: Barnes-Hut t-distributed Stochastic Neighbor Embedding (BH-tSNE) (62) and Uniform Manifold Approximation (UMAP) (63). The product of large-data-mode genome binning totaled 5, 184 parameter combinations per clustering method resulting in 10, 368 jobs per sample (41, 472 total jobs from all four CAMI 2 datasets). All genome binning results were formatted to bioboxes format version 0.9.0 (64) using the ‘autometa-cami-format’ command, then benchmarked using AMBER (56).

**Table 1.** Parameter configurations utilized during parameter sweep benchmarking.

AMBER computes a variety of classification metrics, including measures of cluster purity (precision), completeness (recall) and their harmonic mean (F1 score). To compute these classification scores AMBER applies a mapping strategy where each putative MAG is assigned to a single reference genome, such that the representation of the reference genome is maximized by genome length. Following MAG reference genome assignment, measures of purity and completeness may be determined. Purity represents the ratio of correct assignments (base pairs that overlap with the mapped genome) to incorrect assignments, quantifying the ability of a binner to construct the respective genome without contaminating it with other genomes’ contigs. Completeness indicates the fraction of the genome represented (as measured by the sum of base pairs of the genome). AMBER also determines the adjusted Rand index which is a normalized measure of the Rand index. The Rand index is a clustering measure which compares partitions of base pairs by their membership within the same or disparate genome. True positives (TP) are determined by whether the base pairs within the same genome are grouped in the same MAG. True negatives (TN) are determined by whether the base pairs of different genomes are grouped in separate MAGs. The Rand Index is the sum of these values divided by the sum of all of the pairs. This then undergoes a normalization transformation by accounting for random clustering resulting in the final computed adjusted Rand index (ARI) metric. A perfectly matched genome pair will return a score of one while a reference genome completely split across different MAGs will return zero. A MAG with all its members (i.e. complete) but with other members of a different MAG (i.e. contaminants) is penalized and will be reflected in the ARI score. Conversely (since ARI is a symmetric measure), pure MAGs with unnecessary fragmentation are also penalized.

## RESULTS AND DISCUSSION

### Benchmarks

#### Genome binning

Our primary point of comparison with Autometa 2 genome binning performance was the CAMI 2 binning challenge (2), in which Autometa 1 took part. In that work, Autometa 1 was found to have high average bin purity compared to other pipelines, but it suffered in terms of other metrics, especially ARI (Figure 2), and was judged to have low performance overall. However, as this assessment reflected the default parameters of Autometa 1, we present a more complete exploration of the additional user-defined parameter set here with Autometa 2 (Table 1). Across this parameter sweep, we found that ARI values higher than other binners and often close to the ground truth were achievable, while maintaining binned percentages either comparable to or surpassing other binners (Figure 2). Binned percentage was the primary factor which suffered with worsening metagenomic assembly quality (i.e. gold standard versus MegaHit), but ARI values were roughly comparable between assembly qualities. In Figure 2, the original Autometa 1 benchmarks from the CAMI 2 challenge are distinct from the Autometa 2 parameter sweep results, and this is attributable to the addition of both GC and coverage standard deviation limits in Autometa 2. Our results show that while these limits can lower F1 score and sometimes binned percent, they invariably increase ARI.

**Figure 2.**
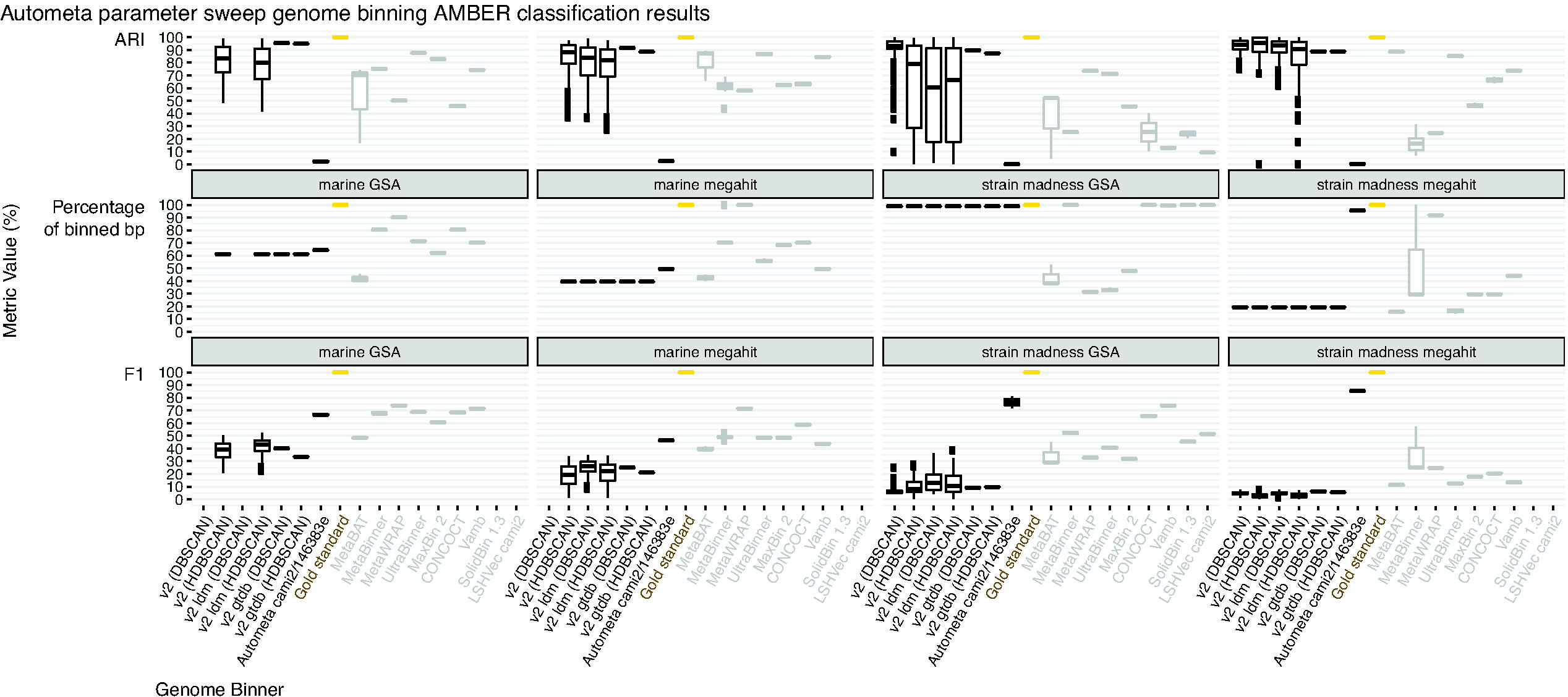
Autometa consistently outperforms all other genome binning tools, reaching near perfect scores according to clustering based on Adjusted Rand Index (ARI), with the tradeoff of lower F1 score. The Autometa 2.1.0 DBSCAN jobs for the marine GSA and megahit datasets did not finish due to memory requirements and are therefore absent.

On the basis of our parameter sweep, it appears Autometa 2 is highly reliable when grouping contigs together (as well as for choosing to leave contigs from disparate genomes separate). In fact, across all four datasets of seq and bp-weighted ARI, Autometa 2 outperformed all other binners, reaching near perfect scores (Figure S4). This tendency towards stringency excludes putative contamination at the expense of leaving genomes incomplete or fragmented. In other words it results in increases in precision, lower misclassification rates and consequently decreases in recall, completeness and F1 score measures.

Understanding Autometa’s genome binning behavior via these metrics provides future algorithmic directions such as implementing a refinement process where high-quality fragmented MAGs could be merged into more complete genomes, suggestive of similar consensus clustering approaches as already outlined (e.g. https://github.com/huangpq2019/ultrabinner and (48)). If implemented correctly, one would expect an improvement in completeness, recall and overall F1 score, while maintaining equivalent performance in precision, purity and misclassification rates. Across all datasets benchmarked, large-data-mode was the majority leader corresponding to ARI, F1 score and average completeness (bp-weighted), while ranking below the autometa-binning entrypoint in average purity (Table S1). Generally the HDBSCAN clustering method occurred more frequently in the best performing parameter configurations for the gold standard assemblies. In contrast, DBSCAN performed better with the megahit assemblies. Interestingly, the highest scoring completeness parameter configurations used relatively low completeness thresholds of 10%, 20%, 30%, and 70% and one purity configuration (10%). This is in opposition to the expected result of higher completeness configurations recovering more complete MAGs. The cutoffs at ten and twenty percent completeness yielded better results in the megahit assemblies whereas the higher thresholds of thirty and seventy percent completeness performed better in the gold standard assemblies (marine and strain madness, respectively). In contrast, the lowest misclassification rates and correspondingly high ARI values (medians ranging from 93-95%) were observed in the results configured with relatively high completeness (e.g. 50%, 80% and 90%) and purity (e.g. 40%, 90%) thresholds.

This behavior is important to consider when choosing what is of interest in a particular dataset. From the parameter sweep approach, it appears Autometa’s overall tendency is to be more stringent rather than lenient, with most parameter sets recovering highly-reliable (yet fragmented) MAGs (Figure S5). Some parameter sets however may degrade the reliability of these MAGs with slight improvements in completeness and F1 score. This is typified by the Autometa 1-like approach of setting limits on the standard deviation of GC and coverage in bins to infinity (Figure S1). The broad distribution displayed by these results highlight the adaptability of Autometa and the scope of fine-tuning available to achieve the desired genome binning result. In settings where a ground truth is not known, different Autometa binning results could be compared with an independent method for estimating genome quality, such as CheckM (65). The effects of Autometa’s user-defined parameters is in contrast to the previously mentioned results from Meyer et al. where parameter changes across all tools accounted for relatively minor changes in performance (approximately 3% as mentioned by Meyer et al.) (2).

#### Taxon binning

To improve Autometa’s performance, we set out to assess the validity of its taxon binning predictions. Robust taxon assignment at each canonical rank may improve genome binning performance in multiple ways. For example, application of a taxon filter to “denoise” a sample. This is particularly useful regarding host-associated metagenomes (14, 22), where any Eukaryotic contamination may be removed. Second, taxon-aware genome binning algorithms may use these taxon bins to confine contigs under consideration for placement within a putative MAG, in order to avoid clustering on very large sets of contigs.

AMBER was used to compute taxon binning metrics (see Genome Binning section for details on metrics) to assess performance of each taxon binner’s results. These metrics were determined across all canonical ranks (kingdom, phylum, class, order, family, genus, species) to demonstrate the scope of each taxon-binner’s capabilities and limitations. Benchmarking was performed using simulated communities (see supplementary Figure S6) and the CAMI 2 datasets, published by Miller et al. and Meyer et al., respectively (2, 3). Autometa consistently outperformed all other taxon-binners (Diamond, Kraken2, MMSeqs2) for the simulated community benchmarking (Figure S6). To more broadly compare Autometa to existing tools, Autometa CAMI 2 taxon binning results were benchmarked against previously published results of Meyer et al. (2) (Figure 3, Table S2).

**Figure 3.**
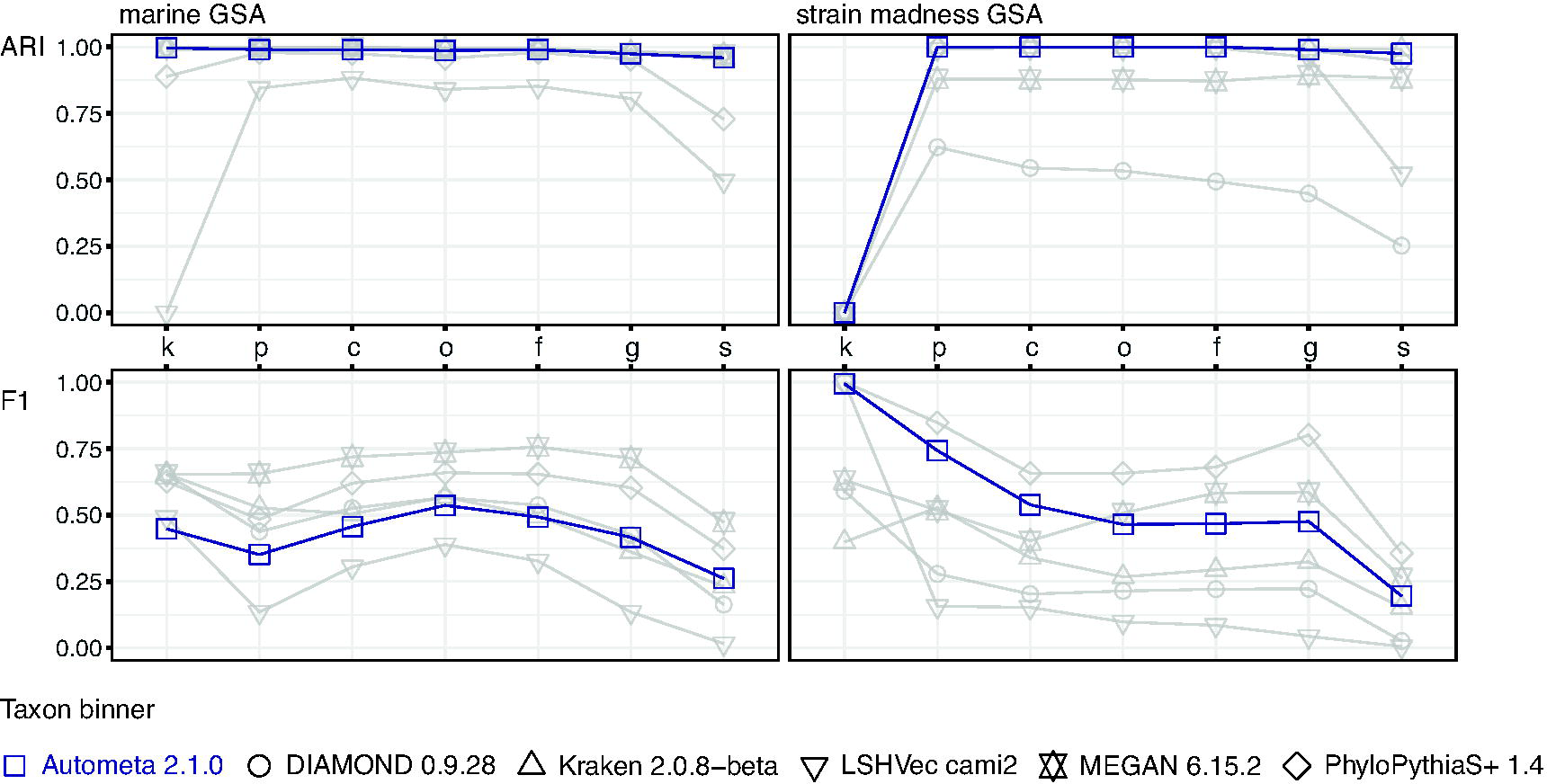
Taxon binning benchmarks against CAMI 2 marine and strain madness gold standard assemblies. Metrics were determined according to canonical ranks, corresponding to kingdom (k), phylum (p), class (c), order (o), family (f), genus (g) and species (s). Autometa again achieves highly reliable results according to the adjusted Rand index (ARI) and according to F1 score is amongst the top three taxon binners as classifications are performed on more specific canonical ranks. Both metrics are bp weighted.

For the strain madness dataset, Autometa is more performant than all except PhyloPithiaS+, according to F1 score, at the kingdom, phylum and class canonical ranks until also being outperformed by MEGAN for the lower ranks (order, family, genus, species). Similarly, Autometa is near-perfect by ARI, with Kraken 2 being slightly better at the species rank. Taking both F1 score and ARI together, Autometa appears to be more consistent at accurately assigning contigs their respective taxon across all canonical ranks (Figure 3). PhyloPithiaS+ may be an optimal choice for replacement of Autometa’s taxon binning sub-workflow when working with datasets containing high amounts of strain overlap (i.e. high levels of microdiversity), due to its consistency across canonical ranks by both F1 and ARI metrics. However, this software package presented difficulties in set up and is no longer maintained, rendering it unsuitable for integration into the Autometa pipeline. In contrast, according to ARI, PhyloPithiaS+ always performed worse than Autometa against the marine dataset with a difference of −0.23 at the species rank. In regards to the marine dataset, it appears MEGAN may be the optimal taxon binning approach. These results suggest that replacing Autometa’s taxon binning results with MEGAN’s for Autometa’s taxon-aware genome binning workflow may improve Autometa’s overall genome binning performance. However, MEGAN is commercial software and consequently makes integration impractical as its requirement would unnecessarily restrict the Autometa user base to MEGAN users.

When considering taxon binner performance collectively with F1 score and ARI across both datasets, Autometa appears to be as reliable as other existing state-of-the-art taxon binning algorithms. Autometa seems to maintain a tendency towards stringency, whereby it is conservative in only assigning contigs when confident about their specific taxon. The trade-off is a result that contains fewer contigs assigned a specific taxon. It should also be noted that across all datasets and all tools, at the species level the maximum F1 score never reached greater than half the theoretical maximum of one. This underscores the overall necessity for further taxon binning method development.

Another aspect of Autometa’s taxon binning pipeline is that it allows deconvolution of sequences derived from different kingdoms, which can be especially important in many environmental and host-associated metagenomes, where either eukaryotic microorganisms or the host genome may interfere with prokaryotic genome binning. Therefore, we reexamined two host-associated metagenomes (10, 22), using our manually curated binning results for two specific uncultured symbionts as the ground truth. We assessed the performance of both Autometa v2 and v1 (as well as v2 under large data mode) against binning pipelines not available at the time of our v1 publication (Figure 4, left). With respect to the “ground truth”, all Autometa versions scored highly as measured by F1, although the Autometa v2 run without large data mode could not achieve the maximum F1 for “*Candidatus* Thermophylae lasonolidus”. This result may be due to the presence of a multi-copy biosynthetic gene cluster that exhibits differing nucleotide frequencies in the genome. MaxBin2 did not achieve F1 scores as good as the best Autometa scores, but VAMB and MetaBat2 approached similar good scores. However, it should be noted that F1 score only reflects precision and recall of a specific ground truth genome, and additional contaminants in the bin would not detract from a good F1 score. Correspondingly, we find that MaxBin2, VAMB and MetaBat2 all suffered from significant contamination from eukaryotic, viral or archaeal contigs, as well as contigs unclassified on the kingdom level (Figure 4, right).

**Figure 4.**
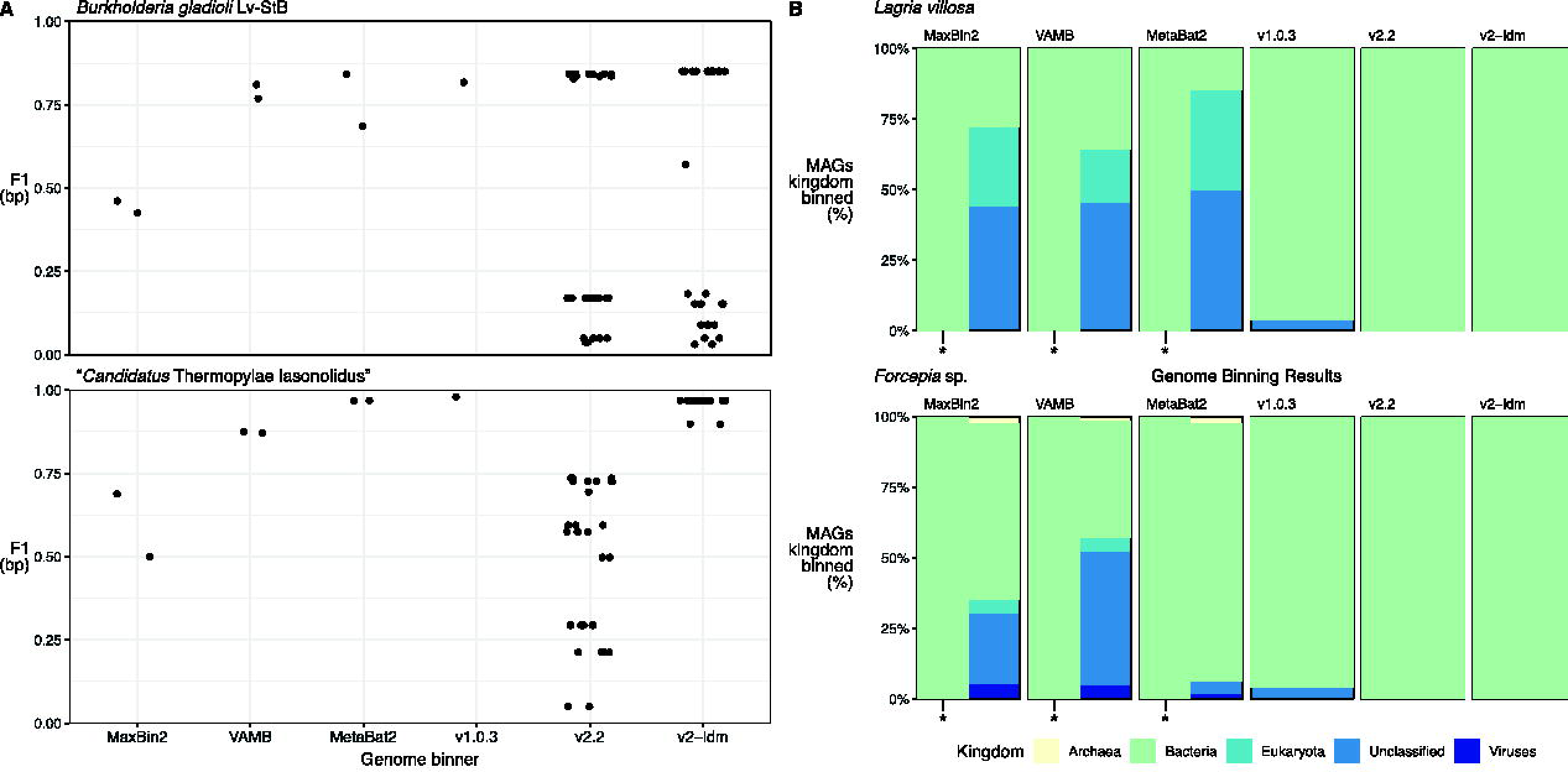
Genome binning results for a beetle (*Lagria villosa*) and sponge (*Forcepia* sp.) metagenome. Panel A displays length-weighted F1 scores for two BGC-containing genomes *Burkholderia gladioli* Lv-StB and *“Candidatus* Thermopylae lasonolidus” pertaining to *L. villosa* and *Forcepia* sp., respectively. Autometa v2 and Autometa large-data-mode (v2-ldm) outperform other genome binners for the *B. gladioli* Lv-StB genome. Autometa v1.0.3 and Autometa large-data-mode (v2-ldm) outperform other genome binners for “*Ca.* Thermopylae lasonolidus”. Panel B depicts the kingdom percentage of MAGs recovered by genome binner for input metagenomes. An (*) indicates genome binning results when using only the bacterial classified sequences as annotated by the Autometa v2 taxon binning workflow. All genome binning results from other genome binners incorporate sequences from other kingdoms, reinforcing the benefits of pre-processing host-associated metagenomes by taxon-binning.

#### Large data mode

In CAMI 2 datasets, large data mode persistently performed comparably to the conventional Autometa pipeline, across the parameter sweep (Figure 2). In both conventional and large data mode, binning of the gold standard marine assembly could not be completed when using DBSCAN as the clustering algorithm, because it requires much more time and memory than HDBSCAN (66). However, even utilizing HDBSCAN for the conventional pipeline, a greater fraction of sample parameter combinations could be completed in large data mode versus the regular approach (Figure 5). In the simulated datasets, there was a tradeoff with large data mode, which exhibited lower F1 scores than the conventional pipeline except in the most complex datasets (Table S3).

**Figure 5.**
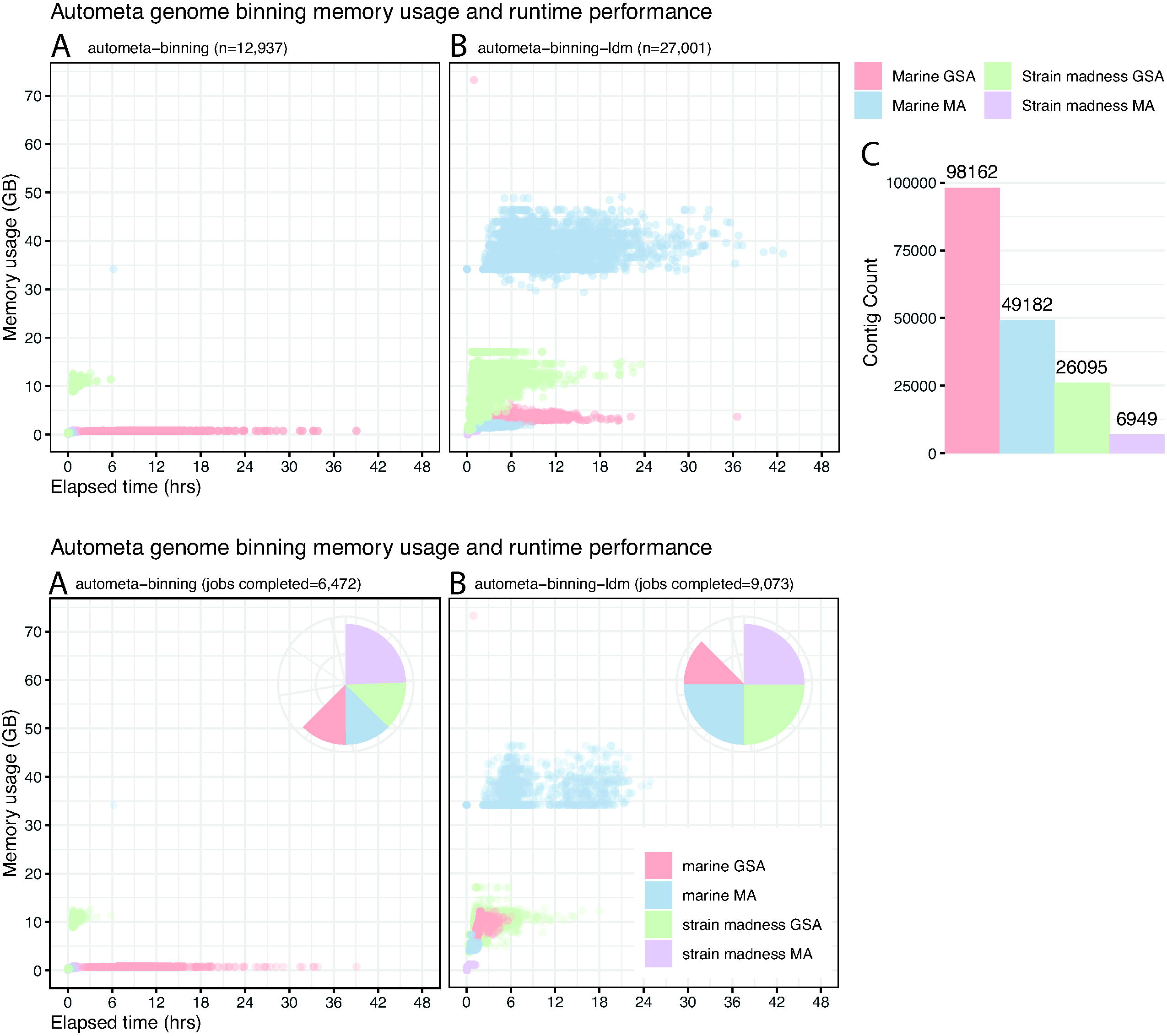
Autometa genome binning runtime and memory usage benchmarks against the CAMI 2 communities. Each point represents a completed genome-binning result. Panel A represents the number of completed runs retrieved using the ‘autometa-binning’ command with 6, 472 total results recovered. Panel B shows completed results using the large-data-mode ‘autometa-binning-ldm’ command which was capable of computing 9, 073 configurations with processing times under 48 hours and RAM usage (mostly) under 50GB. Each entrypoint was provided with the same set of 10, 368 parameter configurations. Point color represents the different source datasets used in testing, and the inset pie charts show the proportion of parameter combinations that were completed within 48 hours for autometa-binning and autometa-binning-ldm.

### Overview of improvements

Autometa has seen widespread use by researchers across the world and along the way its users have provided helpful feedback and suggestions which have guided the second release’s development roadmap. Primary among these difficulties, now amended, is Autometa’s installation. The Autometa library is now distributed via bioconda and docker, allowing the construction of the compute environment and installation to be performed in a few simple commands. This has been complemented with extensive documentation allowing accessible user customization. Efforts have also been made towards making Autometa fault-tolerant, scalable and end-user focused with implementations of nextflow workflows following nf-core standards. These implementations provide access to an easy-to-use graphical user interface (GUI) to aid non-bioinformatics oriented researchers. Likewise, users with access to high-performance compute facilities (or cloud compute infrastructures) may also take advantage of the scalability offered by the Autometa nextflow workflow. Pre-processing, taxon binning, genome binning and binning refinement tasks have been modularized within the Autometa library, providing a structured foundation for future long-term use, maintenance and development. Moreover, Autometa’s “large-data-mode” has been presented in an effort to scale genome binning methods to high-complexity environmental samples. The second major Autometa release provides updates to support multiple areas within metagenome mining and was completely refactored to support straightforward extension and integration with existing ‘omics workflows. These measures were taken to continue to refine Autometa as a key tool for evolutionary studies as well as for aiding culturing and synthetic biology efforts.

### Current challenges

Despite these upgrades, Autometa (as well as other binners) continue to struggle with highly-complex metagenomes where many organisms are novel. During the LCA step of the taxon binning subworkflow, Autometa only uses the hits whose bitscore is within the top 10% of the top hit and is limited to 200 hits total to assign a taxonomic ID, discarding the remaining hit information respective to each query sequence. Furthermore, Autometa fails to account for the divergence of query sequence from its BlastP hits. This may lead to conservative classification of divergent contigs resulting in fewer taxon-assigned contigs as was reflected in the CAMI 2 taxon-binning benchmarks. At this time Autometa does not use all of the information gathered from sequence similarity searches against NCBI’s non-redundant (nr) protein database. Adding to its existing limitations in taxon-binning, draft-quality metagenome-assembled genomes (MAGs) are included in NCBI’s nr database construction. This inclusion introduces the potential for erroneous assignments of subjects to taxids, with these errors being perpetuated by the incorrect genome binning of a contig into a draft-quality MAG. The prevalence and magnitude of this error may merit further investigation. Conversely, these artifacts may resolve themselves through continued revisions of NCBI’s draft-quality MAGs following continued sequencing efforts.

Improving Autometa’s taxonomic classification methods, to deal with database idiosyncrasies and the common presence of novel species, is crucial in order to allow partitioning of large datasets to make the individual fractions more tractable for embedding and clustering algorithms. Algorithmic and computational developments have produced ever-increasing metagenome assembly sizes. As metagenome assembly algorithms scale, preprocessing and genome-binning tasks must similarly progress. Autometa’s available clustering implementations (DBSCAN and HDBSCAN) scale with the input size better than the worst case scenario, but are far from the theoretical best case. In Big O terminology, the computational resources they need scale with the input size, *n*, below O(n^2^) complexity, yet are unable to attain even O(n log (n)) complexity (66). This limitation is partially addressed by reducing the number of contigs to cluster (i.e. limiting *n*) by iterating over taxon binning results rather than the entire metagenome assembly. However, metagenomes with many community members may have an abundance of representatives within each taxon iteration, thereby proportionally deteriorating runtime performance. Thus far, orthogonal metagenome annotations, such as coverage, taxonomy and *k*-mer frequencies, have helped to improve the manner in which to subset the metagenome prior to clustering. Future algorithmic development such as additional sampling heuristics or annotation techniques may ultimately enhance overall genome-binning efficiency.

## DATA AVAILABILITY

The Autometa source code can be found at https://github.com/KwanLab/Autometa and through Figshare (doi: 10.6084/m9.figshare.21944876). The MetaBenchmarks source code can be found at https://github.com/KwanLab/MetaBenchmarks and through FigShare (doi: 10.6084/m9.figshare.21952610). The simulated datasets and ground truths used in benchmarking were deposited to Figshare (doi: 10.6084/m9.figshare.24070359).

## SUPPLEMENTARY DATA

Supplementary Data are available at NAR online.

## AUTHOR CONTRIBUTIONS

Evan R. Rees: Conceptualization, Data curation, Formal analysis, Investigation, Methodology, Project administration, Validation, Visualization, Writing–original draft, Writing–review & editing, Software. Siddharth Uppal: Conceptualization, Data curation, Formal analysis, Investigation, Methodology, Writing–original, Writing–review & editing, Software. Chase M. Clark: Conceptualization, Data curation, Methodology, Writing–original draft, Writing–review & editing, Software. Andrew J. Lail: Data curation, Software. Samantha C. Waterworth: Writing–review & editing, Software. Shane D. Roesemann: Software, Visualization. Kyle A. Wolf: Software. Jason C. Kwan: Conceptualization, Data curation, Methodology, Project administration, Writing–original draft, Writing–review & editing, Software, Supervision, Funding acquisition

## Supporting information

Graphical Abstract

## ACKNOWLEDGEMENTS

The authors would like to thank Brian Couger for his discussions on the development of the large-data-mode algorithm.

## FUNDING

This work was supported by the U.S. National Science Foundation [DBI-1845890 to J.C.K., E.R.R., A.J.L., S.C.W., S.D.R. and K.A.W.]. C.M.C. was supported by an NLM training grant to the Computation and Informatics in Biology and Medicine Training Program [NLM 5T15LM007359]. This research was performed using the compute resources and assistance of the UW-Madison Center For High Throughput Computing (CHTC) in the Department of Computer Sciences. The CHTC is supported by UW-Madison, the Advanced Computing Initiative, the Wisconsin Alumni Research Foundation, the Wisconsin Institutes for Discovery, and the National Science Foundation, and is an active member of the OSG Consortium, which is supported by the National Science Foundation and the U.S. Department of Energy’s Office of Science. Funding for open access charge: National Science Foundation.

## CONFLICT OF INTEREST

The Kwan lab plans to offer their metagenomic binning pipeline Autometa on the paid bioinformatics and computational platform BatchX (https://www.batchx.io) in addition to distributing it through open source channels.

## Notes

https://github.com/KwanLab/Autometa

https://github.com/KwanLab/MetaBenchmarks

https://doi.org/10.6084/m9.figshare.24070359

