## Supplementary figures and images for "Autometa 2: A versatile tool for recovering genomes from highly-complex metagenomic communities"

### Figure S1

Autometa v1 behavior genome binning AMBER classification results

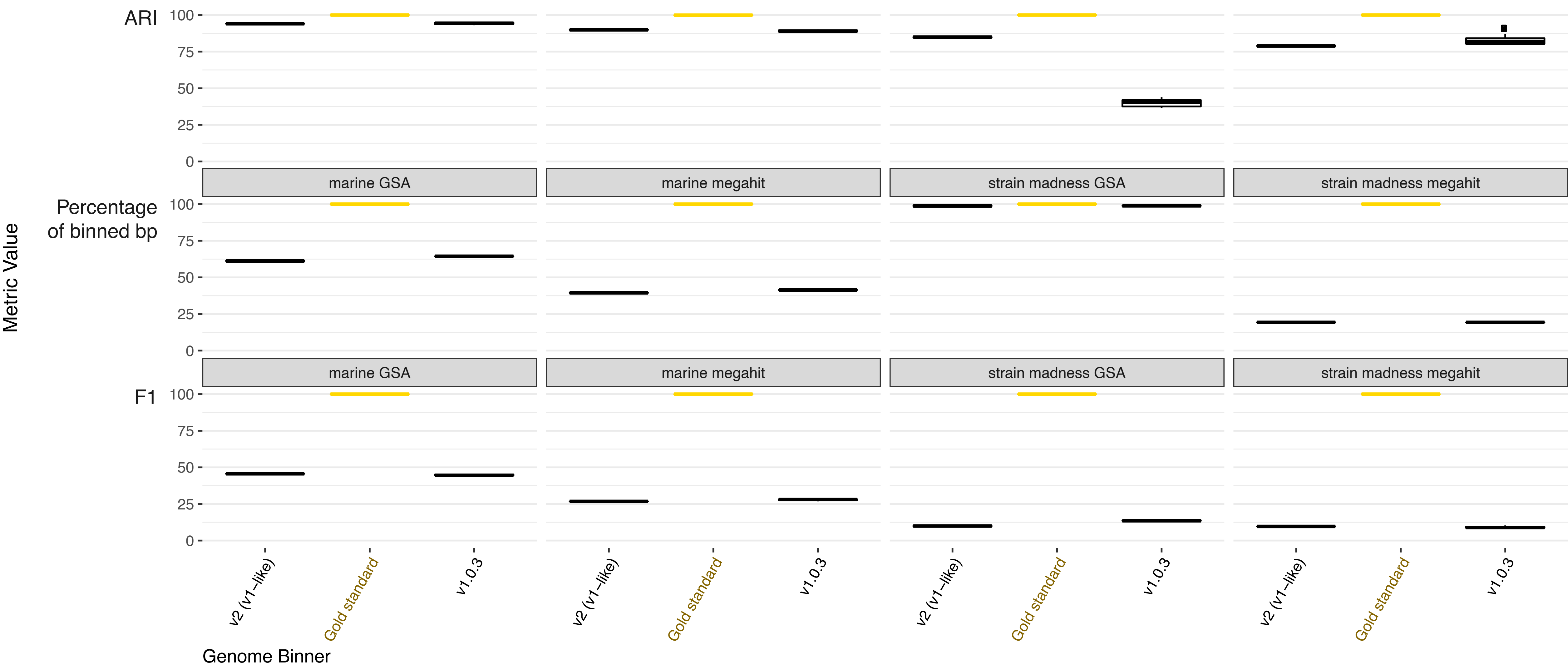

### Figure S4

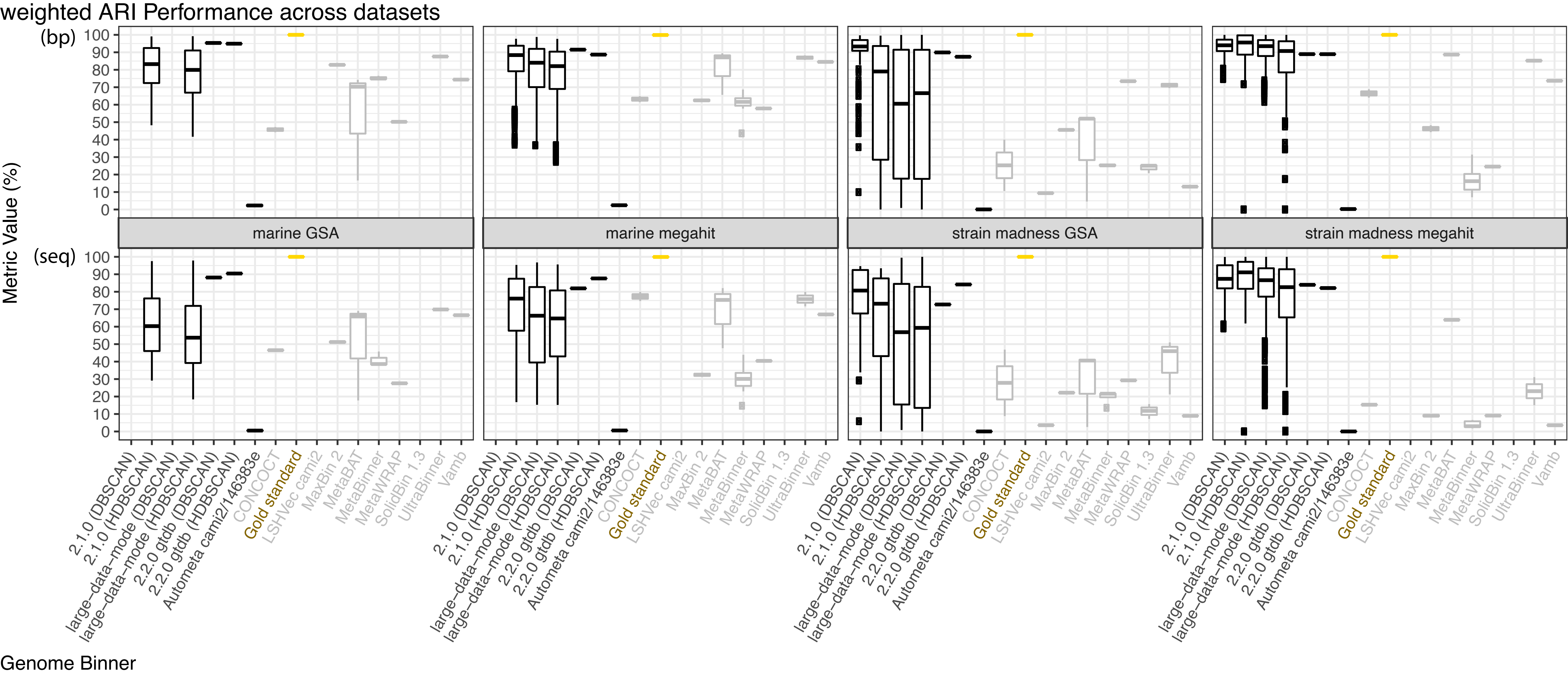

### Figure S5

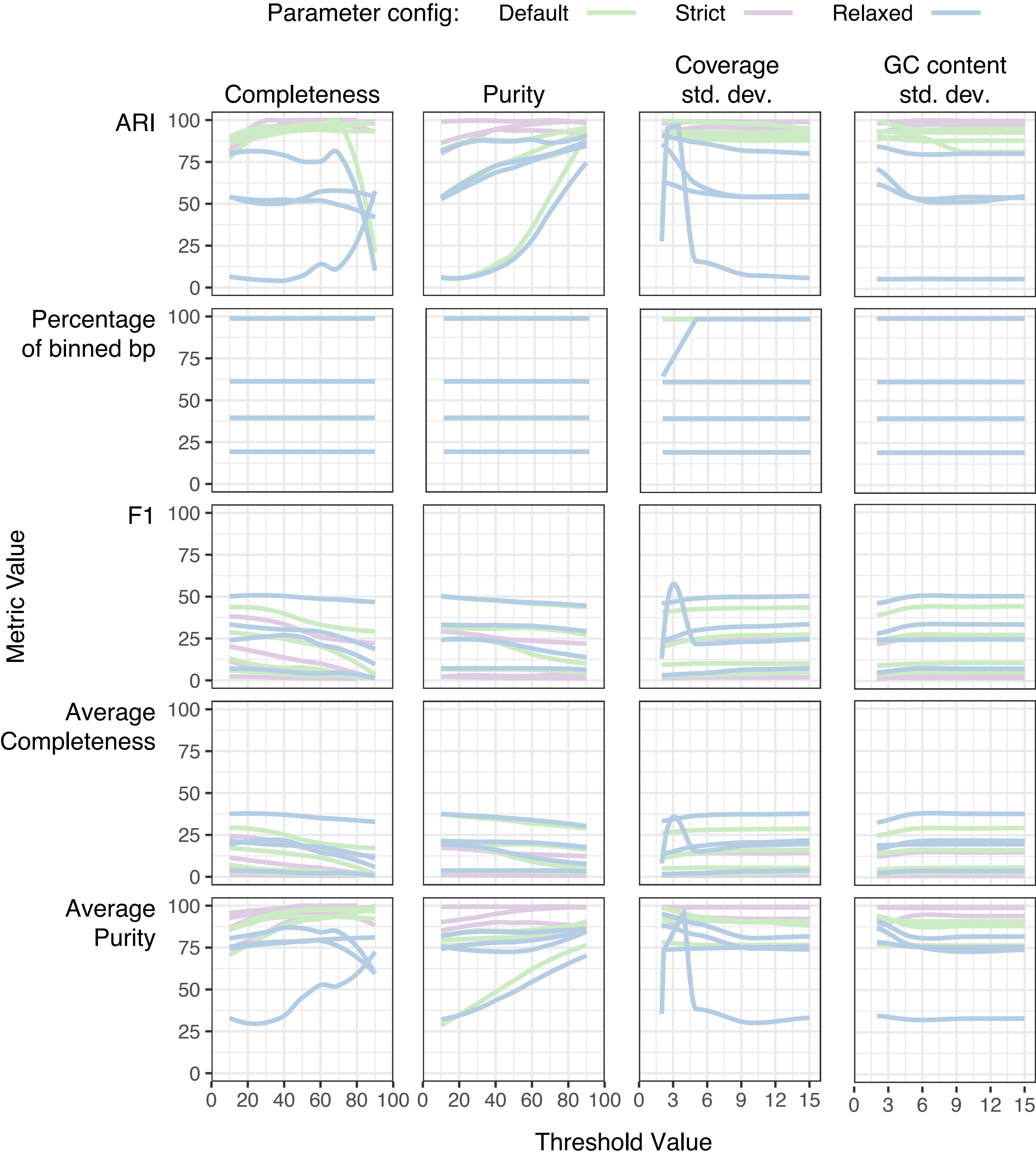

### Figure S6

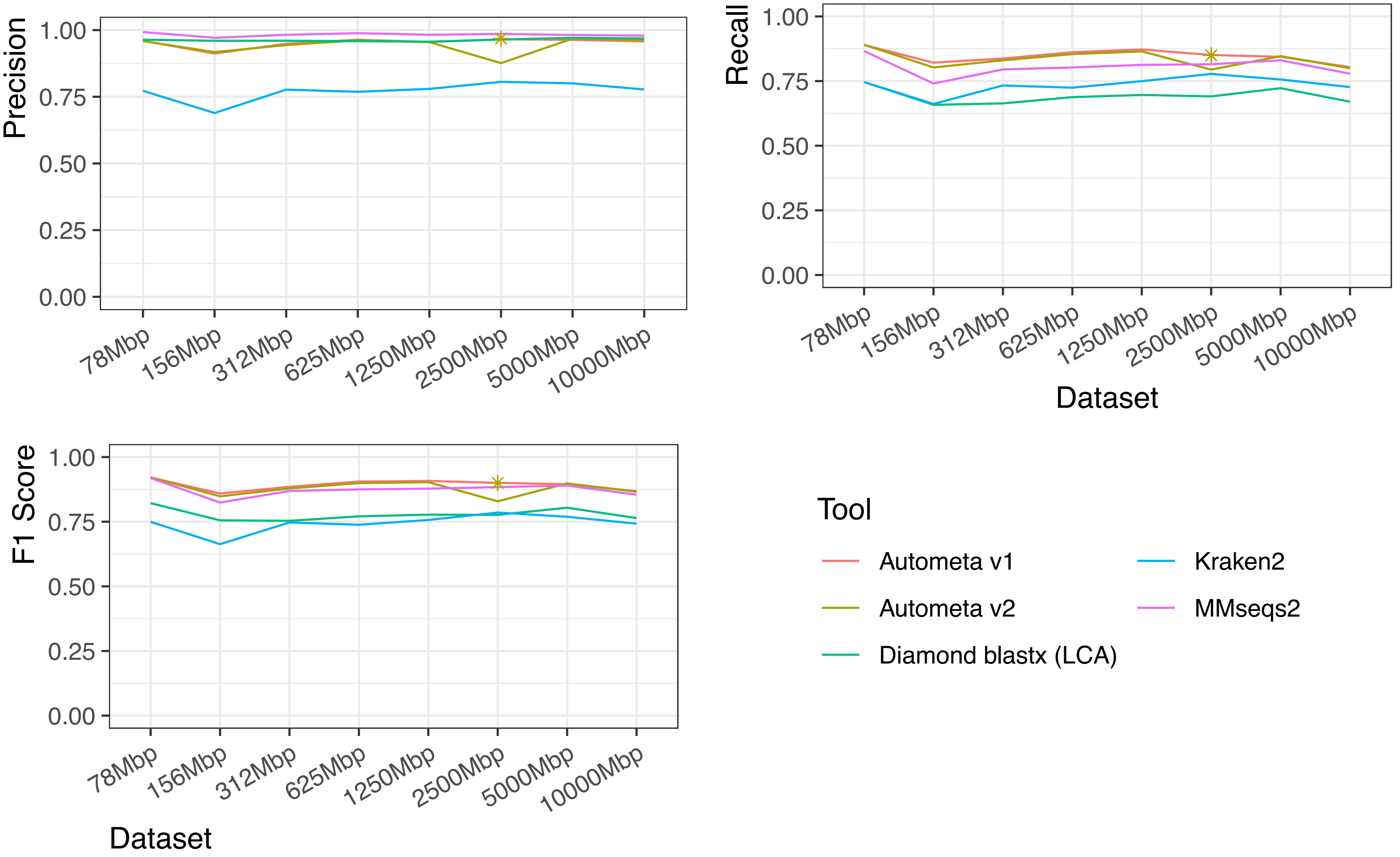

### Graphical Abstract

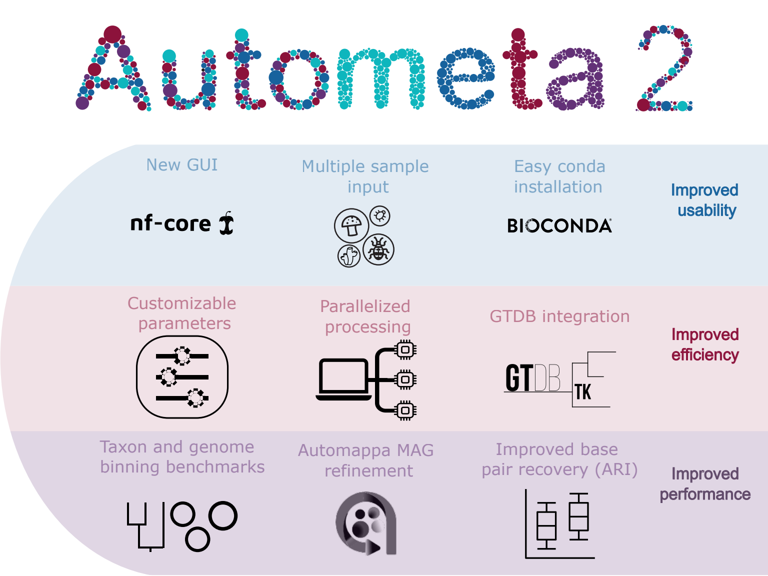
